# Pyroptotic cell-derived factors promote osteoclast differentiation

**DOI:** 10.64898/2026.09.03.748049

**Authors:** Yongjia Li, Chun Wang, Canxin Xu, Wei Zou, Khushpreet Kaur, Ria Rohatgi, Nitin Pokhrel, Nidhi Rohatgi, Yousef Abu-Amer, Gabriel Mbalaviele

## Abstract

The NLRP3 D301N substitution, the murine ortholog of the human D303N variant, causes NLRP3 constitutive activation (NLRP3^CA^) and inflammasome assembly. Here, we show that priming signals induced by lipopolysaccharide (LPS) are sufficient to trigger GSDMD-dependent pyroptosis in NLRP3^CA^-expressing bone marrow-derived macrophages (BMDMs), but not in wild-type (NLRP3^WT^) cells. NLRP3^CA^ mice exhibit elevated IL-1β secretion and LDH release in bone marrow supernatants under both basal and LPS-challenged conditions. Other cytokines (e.g., TNF-α) increased comparably between genotypes, indicating that NLRP3^CA^ specifically amplifies inflammasome-dependent responses. Conditioned medium (CM) from LPS-treated NLRP3^CA^ BMDMs significantly enhanced osteoclast (OC) differentiation *in vivo* and *in vitro* compared to NLRP3^WT^ CM, as did bone marrow supernatants from NLRP3^CA^ mice. This osteoclastogenic activity was largely independent of IL-1β, as demonstrated by experiments using IL-1β-deficient NLRP3^CA^ BMDMs, IL-1β-neutralizing antibodies, and genetic deletion of the IL-1 receptor. The osteoclastogenic factors were primarily soluble proteins, as heat inactivation and proteinase K digestion markedly reduced OC formation. Extracellular vesicles (EVs) from NLRP3^CA^ CM modestly promoted OC differentiation, but EV-depleted supernatants retained full activity, indicating that the primary mediators are soluble proteins not carried within EVs. Together, these findings establish NLRP3^CA^ as a well-controlled model for studying pyroptosis and demonstrate that pyroptotic cells release soluble proteins that drive osteoclastogenesis independently of IL- 1β.

## Introduction

Healthy bone mass is maintained through a dynamic balance between osteoblast-mediated bone formation and osteoclast-mediated bone resorption (1, 2). Inflammation disrupts this balance by favoring osteolysis, thereby contributing to pathological bone destruction in a range of diseases, including osteoporosis, rheumatoid arthritis, periodontal disease, and periprosthetic osteolysis (3–5). Osteoclasts are derived from monocyte/macrophage-lineage precursors and differentiate primarily in response to macrophage colony-stimulating factor (M-CSF) and receptor activator of nuclear factor-κB ligand (RANKL)-RANK signaling, which is negatively regulated by the decoy receptor osteoprotegerin (OPG) (6, 7). Pro-inflammatory cytokines, including TNF-α, IL-1β, IL-6, and IL-17, further promote osteoclastogenesis by increasing RANKL expression and potentiating RANK-mediated signaling cascades that converge on NF-κB, c-Fos, and NFATc1 (8, 9).

Emerging evidence implicates inflammasomes in bone pathologies associated with dysregulated or persistent inflammation (10, 11). Inflammasomes are supramolecular innate immune complexes that activate caspase-1, which catalyzes the maturation of IL-1β and IL-18 and the cleavage of gasdermin D (GSDMD) (12–14). The resulting amino-terminal GSDMD fragments oligomerize and form pores in the plasma membrane, facilitating the release of mature IL-1β and IL-18 and the passage of small molecules (13, 15). Excessive GSDMD-dependent pore formation compromises plasma membrane integrity and promotes membrane rupture through Ninj1, ultimately resulting in pyroptosis, a lytic form of programmed cell death that releases intracellular contents, including factors that can further propagate inflammation (16, 17).

Among the inflammasomes implicated in bone pathophysiology, the NLRP3 inflammasome is the most extensively studied (11, 18–20). NLRP3 inflammasome activation is classically described as a two-step process. The first step, termed priming or licensing, is initiated by pattern-recognition receptors and cytokine receptors and induces transcriptional and post-translational changes that increase NLRP3 abundance and/or competence for activation, as NLRP3 expression and activity are tightly restrained under homeostatic conditions (21, 22). The second activation or assembly step is triggered by cellular stress in response to structurally diverse stimuli, including extracellular ATP, pore-forming toxins, crystalline and soluble substances (21). An important exception to this canonical two-hit model occurs with gain-of-function NLRP3 mutations, in which constitutively active NLRP3 (NLRP3^CA^) can undergo activation with reduced dependence on a conventional second signal and, in some contexts, requires only priming signals (23, 24).

We recently found that systemic administration of lipopolysaccharide (LPS) to mice causes not only inflammation and bone resorption, as expected, but also stimulates periosteal bone formation (25). Mechanistically, we provided that these bone outcomes are caused by pyroptosis of bone marrow cells (25). In the current study, we show that signals derived from bone marrow pyroptotic cells promote OC differentiation.

## Materials and Methods

### Animal and reagents

*Nlrp3^fl(D301N)/+^* mice, kindly provided by H. Hoffman (University of California, San Diego), and *Nlrp3^fl(D301N)/+^*;*LysM-Cre* mice in which NLRP3 is constitutively activated (NLRP3^CA^) have been previously described (13, 26, 27). *Gsdmd* knockout (*Gsdmd^KO^*) mice were kindly provided by Dr. V. M. Dixit (Genentech, South San Francisco, CA), as previously described (28). IL-1β knockout (*Il1b^KO^*) mice and IL-1 receptor knockout (*Il1r^KO^*) were purchased from Jackson Laboratory (Sacramento, CA). All mice were maintained on a C57BL/6J genetic background, and genotyping was performed by polymerase chain reaction (PCR). All animal procedures were conducted under the guidelines and with the approval of the Institutional Animal Care and Use Committee of Washington University School of Medicine in St. Louis, under protocol no. 25-0227. Lipopolysaccharide (LPS) from *E. coli* and proteinase K were purchased from Sigma-Aldrich (St. Louis, MO), and nigericin from Invivogen (San Diego, CA).

### Cell cultures and osteoclast differentiation

Bone marrow-derived macrophages (BMDMs) were obtained by culturing mouse bone marrow cells in medium supplemented with conditioned medium from CMG14-12 fibroblasts (1:10 dilution), which serves as a source of macrophage colony-stimulating factor (M-CSF), according to established protocols (29, 30). Following 4-5 days of culture in 10-cm Petri dishes, nonadherent cells were removed by PBS washing, and adherent BMDMs were collected using trypsin-EDTA. The harvested BMDMs were either maintained in medium containing CMG14-12 conditioned medium (0.2:10) or cultured with CMG14-12 conditioned medium (0.2:10) plus RANKL (50–100 ng/mL) to induce OC differentiation. At the end of the culture period, cells were fixed with 3.7% formaldehyde and 0.1% Triton X-100 for 10 minutes at room temperature and stained with tartrate resistant acid phosphatase (TRAP) solution using the Leukocyte Acid Phosphatase Kit (Sigma-Aldrich, St. Louis, MO) for 30 min at room temperature. TRAP-positive cells containing three or more nuclei were identified as OCs and counted under a light microscope.

### Generation of BMDMs expressing GFP or constitutively activated IKK2 (IKK2^CA^)

IKK2^CA^, GFP were cloned into the pMX retroviral vector fused with either HA or FLAG tag, as previously described (31). To generate retroviral particles, PLAT-E cells, which stably express retroviral packaging genes, were transfected with expression constructs by using TransIT-LT1 transfection reagent (Mirus Bio, Madison, WI) and media was changed after 24 h. Viral supernatants were harvested after 48 h and immediately used to infect freshly isolated bone marrow macrophages in the presence of 4 μg/ml polybrene (MilliporeSigma) for 24 h. Infected cells were then washed and cultured in medium supplemented with a 1:10 dilution of CMG14-12 conditioned medium. Conditioned medium was harvested 48 h later and used for subsequent experiments.

### RT-PCR

Total RNA was extracted with TRIzol reagent (Life Technologies, Carlsbad, CA) and subjected to phenol–chloroform extraction. Following centrifugation at 12,000 × g, the aqueous phase was collected, combined with an equal volume of 70% ethanol, and further purified using the NucleoSpin RNA II Kit (Clontech Laboratories, Mountain View, CA) according to the manufacturer’s instructions. cDNA was synthesized from 1 μg of purified RNA using the cDNA EcoDry Premix Kit. Quantitative PCR was performed with iTaq Universal SYBR Green Supermix (Bio-Rad; Hercules, CA) on an ABI QuantStudio 3 Real-Time PCR System. Cycling conditions were 50°C for 2 min, 95°C for 10 min, followed by 40 cycles of 95°C for 15 s and 60°C for 1 min. Relative transcript abundance was determined using the 2^−ΔCt method. Primer sequences are provided in Table S1.

### Immunoblotting

Whole cell lysates were prepared in RIPA buffer (50 mM Tris, 150 mM NaCl, 1 mM EDTA, 0.5% sodium deoxycholate, 0.1% SDS, and 1% NP-40) containing protease inhibitors (GenDEPOT, Barker, TX). Protein concentrations were quantified using the Bio-Rad Protein Assay. Equal amounts of proteins were separated by 12% SDS–PAGE and transferred to nitrocellulose membranes. Membranes were blocked and incubated overnight at 4°C with primary antibodies, including anti-NLRP3 antibody (1:1000; Adipogen, San Diego, CA), anti-GSDMD antibody (1:1000; Abcam, Cambridge, MA), anti-caspase-1 antibody (1:1000; Abcam, Cambridge, MA), and anti-β-actin antibody (1:2,000; Santa Cruz Biotechnology, Dallas, TX), followed by incubation with IRDye 800 goat anti-mouse or Alexa Fluor 680 goat anti-rabbit secondary antibodies (Thermo Fisher Scientific, Waltham, MA) for 1 h at room temperature. Immunoreactive bands were detected using an Odyssey Infrared Imaging System (LI-COR Biosciences, Lincoln, NE).

### Histology and histomorphometry

Calvariae were fixed in 10% neutral buffered formalin, decalcified in 14% EDTA for 10 days, embedded in paraffin, sectioned at 5 μm thickness, and subsequently stained with either TRAP or H&E. TRAP-stained sections were analyzed using the Bioquant Osteo image analysis system (Bioquant Image Analysis Corporation, Nashville, TN). OCs were identified as TRAP-positive multinucleated cells located along the bone surface. OC number and surface were measured and normalized to bone surface. Results were expressed as OC number per bone surface (N.Oc.N/BS) and OC surface per bone surface (Oc.S/BS).

### LDH and cytokine Assay

Cell death was evaluated by measuring lactate dehydrogenase (LDH) release in the conditioned medium using the LDH Cytotoxicity Detection Kit (Takara, Mountain View, CA). Levels of cytokines and chemokines were quantified using the V-PLEX Plus Proinflamma tory Panel 1 Mouse Kit (Meso Scale Diagnostics, Rockville, MD). Enzyme-linked immunosorbent assay (ELISA) kits were used to measure the levels of IL-1β (eBioscience, San Diego, NY) and IL-18 (R&D Systems, Minneapolis, MN).

### EVs isolation

Extracellular vesicles (EVs) were isolated using either a commercial exosome isolation reagent ExoQuick Exosome Isolation Kit (System Biosciences, Palo Alto, CA) according to the manufacturer’s instructions, or by ultracentrifugation. For the latter, samples were first centrifuged at 10,000 g for 30 minutes, and the resulting supernatant was then centrifuged at 100,000 g for 90 minutes. The supernatants were collected and stored at –80 °C, while the pellets were washed with PBS and centrifuged again at 100,000 g for 70 minutes. The final EV pellets were resuspended in PBS and stored at –80 °C.

### Statistics

Statistical analyses were performed using GraphPad Prism version 9.0 (GraphPad Software). Student’s *t*-test was used for comparisons between two groups. For multiple-group comparisons, one-way or two-way ANOVA followed by Tukey’s post hoc test or two-way ANOVA followed by Dunnett’s post hoc test was applied, as appropriate. Data are representative of 2-3 independent biological replicates.

## Results

### NLRP3^CA^ can assemble a functional inflammasome in the absence of secondary signals

The NLRP3 D301N substitution in mice (the ortholog of the human D303N variant) causes constitutive NLRP3 (NLRP3^CA^) inflammasome assembly (13, 26, 27). In this study, we investigated the effects of inflammasome assembled by WT NLRP3 (NLRP3^WT^) and NLRP3^CA^ on IL-1β and IL-18 secretion, as well as the release of lactate dehydrogenase (LDH), a biomarker of cell lysis, under basal conditions and following stimulation with lipopolysaccharide (LPS), a well-established inducer of priming signals for NLRP3 inflammasome activation (13). LDH and IL-1β levels were higher in the bone marrow supernatants of NLRP3^CA^ mice compared to NLRP3^WT^ counterparts maintained unchallenged (Fig 1A-C) or exposed to LPS (Fig 1D-F). To model this phenotype *in vitro*, we assessed the effects of PBS or LPS on expanded NLRP3^WT^ or NLRP3^CA^ BMDMs. LDH release, IL-1β and IL-18 secretion, as well as NLRP3 expression, GSDMD and caspase-1 processing were comparable between NLRP3^WT^ and NLRP3^CA^ BMDMs in response to PBS treatment (Fig 1G-J). However, upon LPS stimulation, these responses were markedly elevated in NLRP3^CA^ BMDMs, but not in their WT counterparts (Fig 1G-J). The levels of cleaved GSDMD and caspase-1 fragments were remarkably lower at 24 h compared to 5 h post-LPS treatment, likely reflecting pronounced pyroptotic cell death at the later time point. Consistent with this premise and previous studies (13), LPS induced IL-1β secretion and LDH release by NLRP3^CA^ BMDMs, but not by NLRP3^WT^ BMDMs, responses that were abolished in GSDMD deficient BMDMs (Fig S1). We also assessed the expression of several cytokines, including IL- 2, -4, -5, -6, -10, -12, TNF-α, IFN-Ɣ, and CXCL1. The levels of these cytokines, which were similar between NLRP3^WT^ and NLRP3^CA^ BMDMs at baseline, increased comparably between WT and mutant cells in response to LPS stimulation (Fig S2), consistent with the well-established evidence that their expression is not directly dependent on inflammasome activation. Thus, priming signals such as those induced by LPS are sufficient to trigger pyroptosis in cells expressing NLRP3^CA^, but not NLRP3^WT^, providing a well-controlled model for investigating the role of pyroptosis in downstream biological processes.

**Figure 1.**
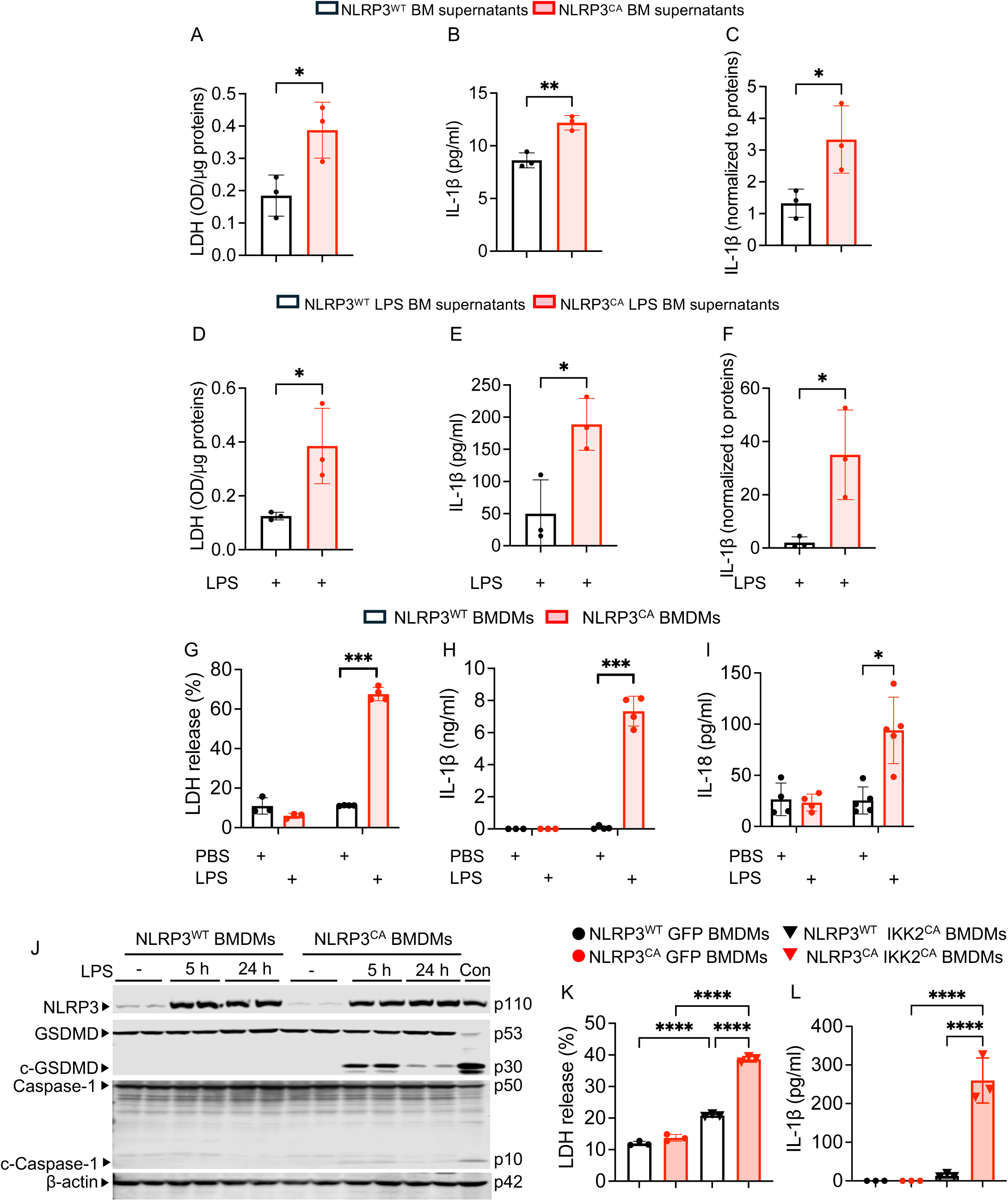
NLRP3CA can assemble a functional inflammasome in the absence of secondary signals. (A-C) whole bone marrow supernatants were collected from the femurs and tibias of NLRP3^WT^ and NLRP3^CA^ mice (n=3). Levels of LDH (A) and IL-1β (B, C). (D-F) NLRP3^WT^ mice and NLRP3^CA^ mice were i.p. injected with 1 mg/kg LPS on days 0 and 4, and mice were sacrificed on day 5. Levels of LDH (D) and IL-1β (E, F) in whole bone marrow supernatants collected from the femurs and tibias of LPS-treated NLRP3^WT^ and NLRP3^CA^ mice (n=3). NLRP3^WT^ and NLRP3^CA^ BMDMs were treated or not with PBS or 30 ng/ml LPS for 24 h (G-J), 5 h or 24 h (J). The supernatants were harvested to measure the levels of LDH (G), IL-1β (H), and IL-18 (I), and cell lysates were used for immunoblotting analysis of the indicated proteins (J). Con, control (LPS + nigericin). NLRP3^WT^ BMDMs treated with 30 ng/ml LPS for 3 h then with nigericin for 45 min served as a positive control) (J). (K, L) NLRP3^WT^ and NLRP3^CA^ BMDMs were transduced with PMX expressing GFP or IKK2^CA^. LDH (K) and IL-1β (L). Samples were run in triplicates and data are presented as mean ± SD. *P<.05, **P<.01, ***P<.001, ****P<.0001; Unpaired t-test; one-way ANOVA test, two-way ANOVA test.

LPS primes the expression of NLRP3 inflammasome components through IKK2-NF-kB cascades (21, 32, 33). Therefore, we tested the ability of lentivirus expressing GFP or constitutively activated IKK2 (IKK2^CA^) to trigger inflammasome responses in NLRP3^WT^ BMDMs (hereafter referred to as NLRP3^WT^ GFP BMDMs and NLRP3^WT^ IKK2^CA^ BMDMs, respectively) or NLRP3^CA^ BMDMs (hereafter referred to as NLRP3^WT^ IKK2^CA^ BMDMs and NLRP3^CA^ IKK2^CA^ BMDMs, respectively). The release of LDH and IL-1β in GFP-expressing cells was comparable between NLRP3^WT^ GFP BMDMs and their NLRP3^CA^ GFP counterparts (Fig 1K, L). Expression of IKK2^CA^ modestly increased LDH and IL-1β release in NLRP3^WT^ BMDMs but induced a robust response in NLRP3^CA^ BMDMs compared to the other genotypes (Fig 1K, L). Thus, consistent with previous reports (13), LPS alone is sufficient to trigger inflammasome formation and downstream signaling *in vitro* in BMDMs expressing NLRP3^CA^ but not NLRP3^WT^. Accordingly, because LPS activates IKK2, forced expression of IKK2^CA^ models the effects LPS on NLRP3 inflammasome signaling.

### Pyroptotic cells release factors that promote OC differentiation

To determine the effects of pyroptotic cell-derived factors on osteoclastogenesis *in vivo*, we injected conditioned medium (CM) collected from LPS-treated NLRP3^WT^ BMDMs (NLRP3^WT^ LPS CM) or LPS-treated NLRP3^CA^-expressing BMDMs (NLRP3^CA^ LPS CM) into the calvariae of WT mice. OC number and surface area did not differ significantly between untreated mice and mice treated with NLRP3^WT^ LPS CM (Fig 2A-D). In contrast, both OC parameters were significantly increased in mice treated with NLRP3^CA^ LPS CM compared to those treated with NLRP3^WT^ LPS CM (Fig 2A-D). We further assessed the osteoclastogenic activity of bone marrow-derived soluble factors, collected from bone marrow (BM) supernatants of untreated NLRP3^WT^ and NLRP3^CA^ mice (hereafter referred to as NLRP3^WT^ BM supernatants and NLRP3^CA^ BM supernatants, respectively), as well as from LPS-treated NLRP3^WT^ and NLRP3^CA^ mice (NLRP3^WT^ LPS BM supernatants and NLRP3^CA^ LPS BM supernatants, respectively). The supernatants were added to WT BMDMs (hereafter simply referred to as BMDMs) primed with RANKL for 2 days. NLRP3^WT^ BM supernatants promoted OC differentiation compared with untreated control cultures, although the effect was significantly weaker than that induced by NLRP3^CA^ BM supernatants (Fig 2E-H). Similarly, NLRP3^WT^ LPS BM supernatants enhanced OC differentiation, but to a lesser extent than NLRP3^CA^ LPS BM supernatants (Fig 2I-L).

**Figure 2.**
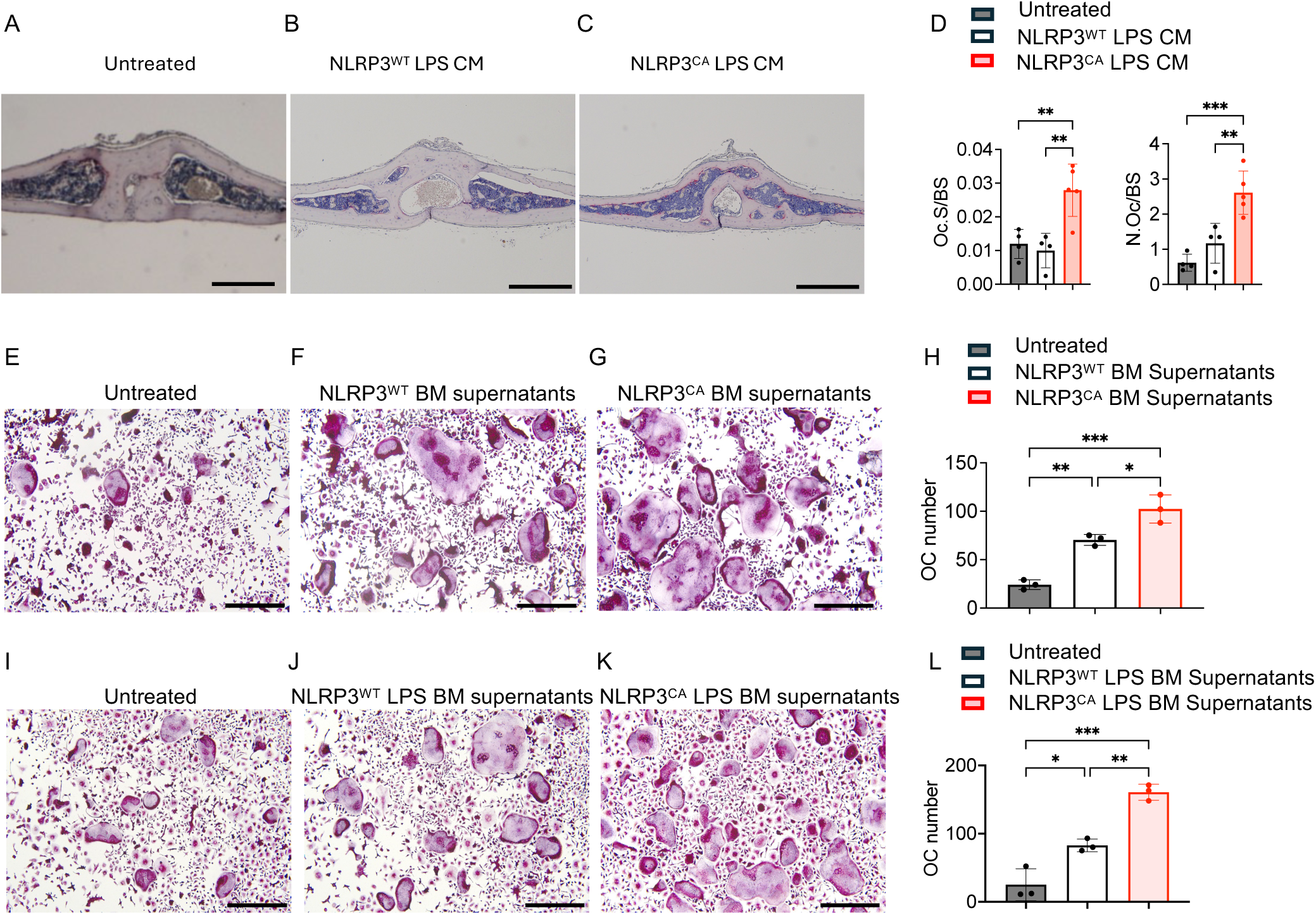
Pyroptotic cell-released factors that promote OC differentiation *in vivo*. (A) untreated mice. (B-D) Conditioned media (CM) from NLRP3^WT^ and NLRP3^CA^ BMDMs treated with 30 ng/ml LPS for 24 h, NLRP3^WT^ LPS CM and NLRP3^CA^ LPS CM, respectively, were injected (50 μl) to WT mice once daily on days 1-4 (n=4-5 mice/group). On day 5, mice were euthanized, and calvariae were harvested TRAP staining of longitudinal sections. Representative TRAP- stained specimens (A-C) and quantitative analysis (D). Scale bar, 450 µm. N.Oc/BS, OC number/bone surface; Oc.S/BS, OC surface/bone surface. (E-H) Whole bone marrow supernatants collected from femurs and tibias of NLRP3^WT^ (NLRP3^WT^ BM supernatants) and NLRP3^CA^ mice (NLRP3^CA^ BM supernatants) and were added to WT BMDMs primed with RANKL for 2 days. Cells were cultured for additional 2-3 days in the presence of RANKL and the supernatants refreshed every day, then were stained for TRAP activity. (E-H) Representative images. Scale bar, 200 µm. The number of TRAP^+^ cells with at least 3 nuclei (OCs) was counted (G). (I-L) NLRP3^WT^ mice and NLRP3^CA^ mice were i.p. injected with 1 mg/kg LPS on days 0 and 4, were sacrificed on day 5, and bone marrow supernatant was collected from femurs and tibias, NLRP3^WT^ BM supernatants and NLRP3^CA^ supernatants, respectively. The supernatants were added to WT BMDMs primed with RANKL for 2 days. Cells were cultured for additional 2-3 days in the presence of RANKL and the supernatants refreshed every day, then were stained for TRAP activity. (I-K) Representative images. Scale bar, 200 µm. The number of OCs was counted (L). Samples were run in triplicates and data are presented as mean ± SD. *P<.05, **P<.01, ***P<.001; one-way ANOVA test.

To model BM supernatant-mediated responses *in vitro*, we evaluated the effects of conditioned media (CM) from PBS- or LPS-treated NLRP3^WT^ or NLRP3^CA^ BMDMs. Specifically, CM from NLRP3^WT^ or NLRP3^CA^ BMDMs treated with PBS (NLRP3^WT^ PBS CM and NLRP3^CA^ PBS CM, respectively) alongside CM from NLRP3^WT^ or NLRP3^CA^ BMDMs treated with LPS (NLRP3^WT^ LPS CM and NLRP3^CA^ LPS CM, respectively) were added to BMDMs primed with RANKL for 2 days. OC differentiation was comparable between WT BMDMs left untreated or treated with NLRP3^WT^ PBS CM, NLRP3^CA^ PBS CM, or NLRP3^WT^ LPS CM (Fig 3A-D, F). By contrast, this response was significantly enhanced in BMDMs treated with NLRP3^CA^ LPS CM (Fig 3E, F). Accordingly, the expression of several OC markers induced by RANKL was further increased by NLRP3^CA^ LPS CM (Fig S3). We next investigated OC differentiation of RANKL-primed BMDMs exposed to CM from GFP- or IKK2^CA^-transduced NLRP3^WT^ (NLRP3^WT^ GFP CM and NLRP3^CA^ GFP CM, respectively) or NLRP3^CA^ BMDMs (NLRP3^WT^ IKK2^CA^ CM and NLRP3^CA^ IKK2^CA^ CM, respectively). OC differentiation was induced to a comparable extent by NLRP3^WT^ GFP CM and NLRP3^CA^ GFP CM, modestly enhanced by NLRP3^WT^ IKK2^CA^ CM, and robustly stimulated by NLRP3^CA^ IKK2^CA^ CM (Fig 3G-L). Collectively, these findings demonstrate that pyroptotic cell-derived factors promote OC differentiation both *in vivo* and *in vitro*.

**Figure 3.**
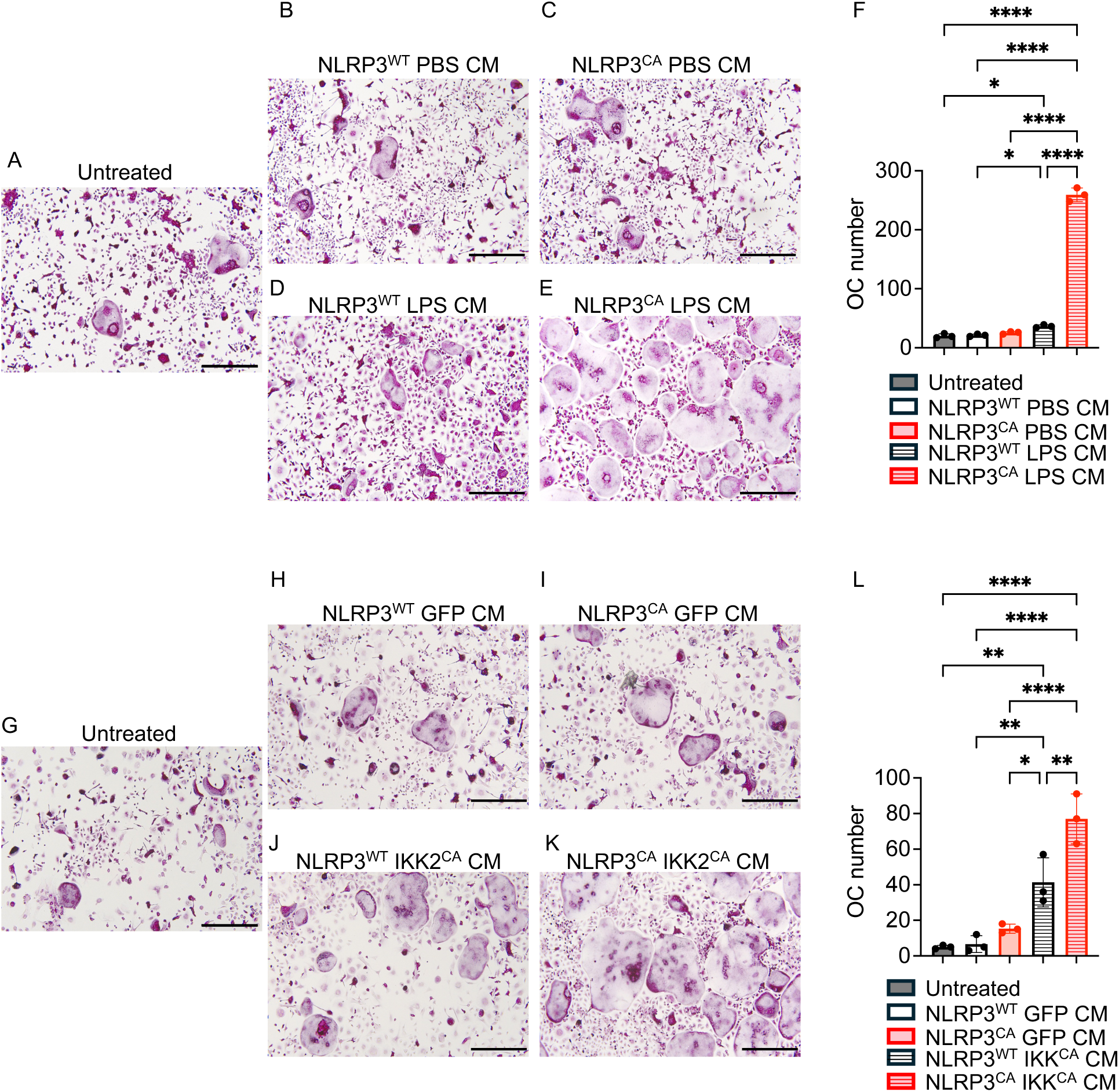
Pyroptotic cell-derived factors promote OC differentiation *in vitro*. (A-F) CM from NLRP3^WT^ or NLRP3^CA^ BMDMs treated with PBS (NLRP3^WT^ PBS CM and NLRP3^CA^ PBS CM, respectively) alongside CM from NLRP3^WT^ or NLRP3^CA^ BMDMs treated with 30 ng/ml LPS for 24 h (NLRP3^WT^ LPS CM and NLRP3^CA^ LPS CM, respectively) were added to BMDMs primed with RANKL for 2 days. Cells were cultured for additional 2-3 days in the presence of RANKL and CM refreshed every day, then were stained for TRAP activity. (A-E) Representative images. Scale bar, 200 µm. The number of OCs was counted (F). (G-L) CM from GFP- or IKK2^CA^-transduced NLRP3^WT^ (NLRP3^WT^ GFP CM and NLRP3^CA^ GFP CM, respectively) or NLRP3^CA^ BMDMs (NLRP3^WT^ IKK2^CA^ CM and NLRP3^CA^ IKK2^CA^ CM, respectively) were added to BMDMs primed with RANKL for 2 days. Cells were cultured for additional 2-3 days in the presence of RANKL and CM refreshed every day, then were stained for TRAP activity. (G-K) Representative images. Scale bar, 200 µm. The number of OCs was counted (L). Samples were run in triplicates and data are presented as mean ± SD. *P<.05, **P<.01, ***P<.001, ****P<.0001; one-way ANOVA test.

### OC differentiation promoted by pyroptotic cell-derived factors occurs largely independently of released IL-1β

NLRP3^CA^ BMDMs secreted approximately 7 ng/mL IL-1β and 100 pg/mL IL-18 in response to LPS (Fig 1H, I). To determine whether these cytokines cooperate with RANKL to regulate OC differentiation, RANKL-primed BMDMs were treated with increasing concentrations of IL-1β or IL-18. IL-1β significantly promoted OC formation at concentrations >5 ng/mL (Fig S4A, B), whereas IL-18 had no effect (Fig S4C, D). To assess the contribution of IL-1β to the osteoclastogenic activity of pyroptotic CM, BMDMs expressing NLRP3^WT^ or NLRP3^CA^, as well as NLRP3^CA^ BMDMs lacking IL-1β (NLRP3^CA^;IL-1β^KO^ BMDMs) were treated with PBS or LPS, and IL-1β secretion and LDH release were measured. PBS-treated BMDMs of all genotypes and LPS-treated NLRP3^WT^ BMDMs released similarly low levels of IL-1β and LDH (Fig 4A, B). LPS induced comparable LDH release in both NLRP3^CA^ and NLRP3^CA^;IL-1β^KO^ BMDMs, indicating equivalent pyroptosis, whereas IL-1β secretion was detected only in NLRP3^CA^ BMDMs (Fig 4A, B). We next examined the effects of CM from these BMDMs on OC differentiation of WT BMDMs. LPS-CM from NLRP3^CA^ BMDMs robustly stimulated OC formation, whereas CM from NLRP3^CA^;IL-1β^KO^ BMDMs produced only a modest reduction in osteoclastogenesis (Fig 4C-I). Consistent with these findings, neutralizing IL-1β antibodies only slightly attenuated the osteoclastogenic activity of NLRP3^CA^ LPS CM, which remained significantly greater than that of NLRP3^WT^ LPS CM regardless of IgG or IL-1β antibody treatment (Fig 4J-L). Furthermore, genetic deletion of the IL-1 receptor (IL-1R^KO^), which abolishes signaling by both IL-1α and IL-1β, did not eliminate OC differentiation induced by NLRP3^CA^ LPS CM compared with NLRP3^WT^ LPS CM (Fig 4M-R). Collectively, these findings indicate that although IL-1β contributes to pyroptosis-induced osteoclastogenesis, it is not the primary mediator of the osteoclastogenic activity present in pyroptotic CM.

**Figure 4.**
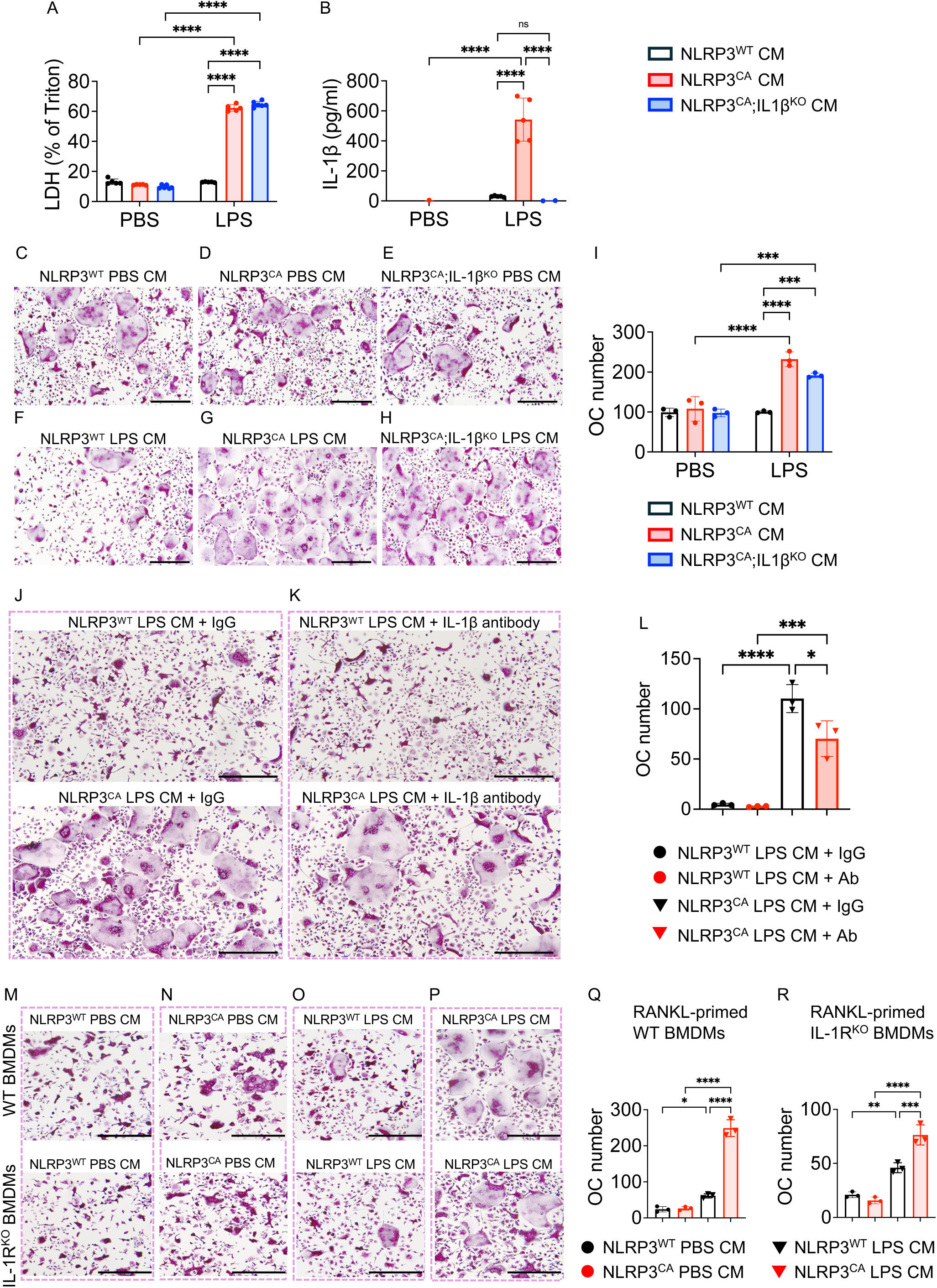
Pyroptotic cell-derived factors promote OC differentiation largely independently of released IL-1β. (A-I) BMDMs obtained from NLRP3^WT^, NLRP3^CA^ and NLRP3^CA^; IL-1βKO mice were treated with PBS or 30 ng/ml LPS for 24 h; CM were collected as NLRP3^WT^ PBS CM, NLRP3^CA^ PBS CM, NLRP3^CA^;IL-1βKO PBS CM, NLRP3^WT^ LPS CM, NLRP3^CA^ LPS CM, and NLRP3^CA^;IL-1βKO LPS CM. The levels of LDH (A) and IL-1 β (B) were measured in CM. CM were added to WT BMDMs primed with RANKL for 2 days. Cells were cultured for additional 2-3 days in the presence of RANKL and CM refreshed every day, then were stained for TRAP activity. (C-H) Representative images. The number of OCs was counted (I). (J, K) Representative images. The number of OCs was counted (L). (J-L) NLRP3^WT^ LPS CM and NLRP3^CA^ LPS CM was added to WT BMDMs primed with RANKL for 2 days in the presence of 10 μg/ml IgG or 10 μg/ml IL-1β neutralizing antibody (Ab). Cells were cultured for additional 2-3 days in the presence of RANKL and CM with IgG or antibody refreshed every day, then were stained for TRAP activity. (M-R) NLRP3^WT^ PBS CM, NLRP3^CA^ PBS CM, NLRP3^WT^ LPS CM and NLRP3^CA^ LPS CM were added to WT and IL-1β^KO^ BMDMs primed with RANKL for 2 days. Cells were cultured for additional 2-3 days in the presence of RANKL and CM refreshed every day, then were stained for TRAP activity. Scale bar, 200 µm. Samples are presented as mean ± SD. *P<.05, **P<.01, ***P<.001, ****P<.0001; one- way ANOVA test, two-way ANOVA test.

### OC differentiation promoted by pyroptotic cells occurs largely through the action of released proteins, some of which are carried within extracellular vesicles

Pyroptotic cells release a variety of molecules, including lipids, metabolites, and proteins (34, 35). To investigate the nature of the osteoclastogenic factors present in the CM, it was left untreated, heat-inactivated, or heat-inactivated followed by proteinase K digestion. As expected, proteinase K digestion altered both the abundance and migration of proteins, whereas heat inactivation had no noticeable effect (Fig S5). Untreated NLRP3^CA^ LPS CM significantly enhanced OC differentiation compared to untreated NLRP3^WT^ LPS CM (Fig 5A, B). This effect was markedly reduced by heat inactivation and further suppressed by combined heat inactivation and proteinase K treatment (Fig 5A, B). These findings suggest that protein components released by pyroptotic cells are critical mediators of osteoclastogenesis.

**Figure 5.**
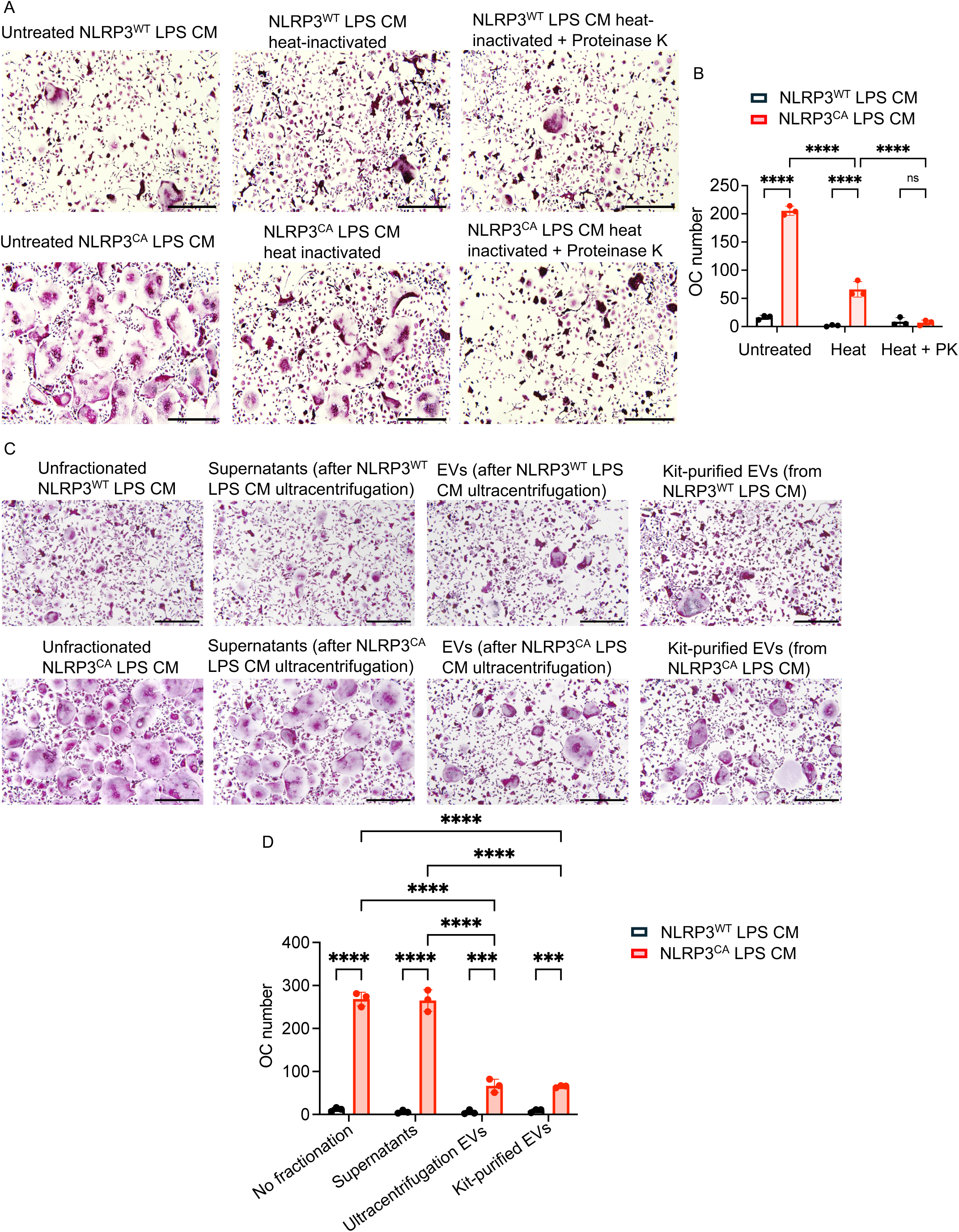
OC differentiation promoted by pyroptotic cells occurs largely through the action of released proteins, some of which are carried within extracellular vesicles. (A-B) NLRP3^WT^ LPS CM and NLRP3^CA^ LPS CM were either heated at 95°C for 30 min alone or heated at 95°C for 30 min followed by treatment with proteinase K (PK) at a final concentration of 100 µg/ml. CM was added to WT BMDMs primed with RANKL for 2 days. Cells were cultured for additional 2-3 days in the presence of RANKL and CM refreshed every day, then were stained for TRAP activity. (A) Representative images. The number of OCs was counted (B). (C-D) BMDMs were treated with unfractionated CM, supernatants (after LPS CM ultracentrifugation) EVs (after ultracentrifugation or kit-purification of LPS CM). CM was added to WT BMDMs primed with RANKL for 2 days. Cells were cultured for additional 2-3 days in the presence of RANKL and CM refreshed every day, then were stained for TRAP activity. (C) Representative images. The number of OCs was counted (D). Scale bar, 200 µm. Samples were run in triplicates and data are presented as mean ± SD. ***P<.001, ****P<.0001, two-way ANOVA test.

A recent study suggested that gasdermin pores can be transported via extracellular vesicles (EVs), thereby propagating pyroptosis to bystander cells (36). To assess whether EVs derived from pyroptotic cells contribute to osteoclastogenesis, BMDMs were treated with unfractionated CM, EVs, or EV-depleted supernatants. EVs isolated from NLRP3^CA^ LPS CM significantly promoted OC differentiation compared to NLRP3^WT^ LPS CM (Fig 5C, D). However, their effect was less pronounced than that of EV-depleted supernatants, which was comparable to unfractionated NLRP3^CA^ LPS CM (Fig. 5C, D). These findings suggest that osteoclastogenic activity is primarily driven by soluble factors released from pyroptotic cells that are not associated with EVs.

## Discussion

Our findings demonstrate that NLRP3^CA^ substantially lowers the threshold for inflammasome signaling and pyroptosis in BMDMs. *In vivo*, NLRP3^CA^ mice exhibit increased basal and LPS-induced IL-1β and LDH release, whereas *in vitro*, LPS is sufficient to induce caspase-1 and GSDMD processing, cytokine secretion, and pyroptotic cell death in NLRP3^CA^ but not NLRP3^WT^ BMDMs. This is consistent with previous studies showing that the NLRP3 D301N mutation allows inflammasome activation in the absence of a conventional secondary signal (13, 26, 27). Importantly, the comparable production of other LPS-induced cytokines between genotypes indicates that the enhanced pyroptosis-dependent responses are not attributable to generalized hyperresponsiveness to LPS but rather reflects selective amplification of the NLRP3-caspase-1- GSDMD axis. The ability of constitutively active IKK2 to reproduce this phenotype further supports the concept that an otherwise canonical priming pathway can become sufficient to drive inflammasome activation when NLRP3 is constitutively active.

The functional consequences of this enhanced inflammasome activity extend beyond the pyroptotic cells themselves. CM from NLRP3^CA^ pyroptotic macrophages consistently promotes OC differentiation more strongly than media from NLRP3^WT^ cells, both following direct administration *in vivo* and in RANKL-primed cultures *in vitro*. These observations are consistent with a growing body of literature linking inflammasome activation and pyroptosis to pathological osteoclastogenesis and bone loss (27, 37). Notably, osteoclastogenic activity is also evident in bone marrow supernatants from unchallenged NLRP3^CA^ mice, suggesting that constitutive inflammasome activation may establish an osteoclastogenic inflammatory milieu *in vivo*. Our findings with NLRP3^CA^ contrast with our previous study, in which a low dose of LPS induced bone marrow cell pyroptosis in WT mice predominantly through the noncanonical inflammasome pathway (25). By contrast, the present findings In the NLRP3^CA^ model implicate canonical NLRP3 inflammasome activation as the principal driver of pyroptosis. Whether these differences reflect the distinct mouse strains, LPS dosing regimens, or other experimental variables remains unclear and warrants further investigation. Nevertheless, the findings from both models support the broader concept that pyroptosis may serve as an important mechanism for transmitting inflammatory and osteoclastogenic signals from myeloid cells to OC precursors.

Although IL-1β is a well-established promoter of OC differentiation, our results indicate that it is not the principal mediator of the osteoclastogenic activity associated with NLRP3^CA^-derived pyroptotic CM. Exogenous IL-1β promotes osteoclastogenesis only at relatively high concentrations, and genetic deletion or neutralization of IL-1β produced only a modest reduction in the activity of NLRP3^CA^ CM. Moreover, the persistence of enhanced osteoclastogenesis in IL- 1R-deficient cultures argues against a dominant requirement for signaling through either IL-1α or IL-1β. This finding is particularly important because previous studies have largely emphasized IL-1β as the major link between inflammasome activation, pyroptosis, and osteoclastogenesis (11, 20, 38, 39). Our results instead suggest that pyroptotic cells release a broader osteoclastogenic secretome capable of promoting RANKL-driven differentiation independently of canonical IL-1 signaling.

Although lipids and metabolites have been shown to mediate biological effects of pyroptotic cells (34), the marked sensitivity of osteoclastogenic activity to heat treatment and proteinase K digestion in the present study supports a predominant contribution from extracellular proteins. Consistent with this interpretation, our previous mass spectrometry analysis demonstrated that LPS-induced pyroptosis of bone marrow cells was accompanied by abundant release of intracellular proteins, and we further showed that the osteogenic activity of pyroptotic cell supernatants was primarily mediated by proteins (25). Although extracellular vesicles derived from NLRP3^CA^ pyroptotic cells retained osteoclastogenic activity, EV-depleted supernatants were substantially more potent, indicating that the major activity resides in soluble, non-EV-associated factors. This finding expands the current concept of pyroptosis from a mechanism primarily associated with IL-1β/IL-18 release to one involving the broader dissemination of bioactive extracellular factors. Together, these data support a model in which constitutive NLRP3 activation enhances pyroptosis and generates a protein-rich extracellular milieu that potentiates RANKL-dependent osteoclastogenesis, with IL-1β contributing only partially to this response.

The limitations of our study include the potential confounding effects of LPS, although LPS was present in cultures of both NLRP3^WT^ and NLRP3^CA^ cells, and its effects on NLRP3 inflammasome signaling were recapitulated in the absence of LPS by forced expression of IKK2^CA^. In addition, the specific pyroptotic factors that promote osteoclast formation were not identified. Despite these limitations, our findings collectively identify pyroptotic macrophages as active regulators of the osteoclastogenic microenvironment and provide a mechanistic link between pathological NLRP3 activation and enhanced bone resorption. This interpretation is consistent with previous work demonstrating that myeloid NLRP3^D301N^ activation promotes osteoclast formation and bone loss, while extending those observations by suggesting that factors released from pyroptotic cells, rather than IL-1β alone, may mediate much of the osteoclastogenic effect.

## Acknowledgements

This work was funded by the National Institutes of Health grants R01 AG077732 and R01 AI161022 to GM. YA was supported by R01 AR082192 and R01 AR081270. The Histology Core was supported by Washington University Musculoskeletal Research Center (NIH P30 AR074992).

Yongjia Li^1^, Chun Wang^1^, Canxin Xu^2^, Wei Zou^1^, Khushpreet Kaur^1^, Ria Rohatgi^1^, Nitin Pokhrel^1^, Nidhi Rohatgi^1^, Yousef Abu-Amer^3^

## Author contributions

Conceptualization: YL, GM

Methodology: YL, CW, CX, WZ, KK, RR, NP, NR

Funding acquisition: YA, GM Supervision: GM

Writing – original draft: YL, GM

Writing – review & editing: YL, CW, YA, GM

## References

1. Bolamperti, S., I. Villa, and A. Rubinacci. 2022. Bone remodeling: an operational process ensuring survival and bone mechanical competence. Bone Res. 10: 48.

2. Sims, N. A., and T. J. Martin. 2020. Osteoclasts Provide Coupling Signals to Osteoblast Lineage Cells Through Multiple Mechanisms. Annu. Rev. Physiol. 82: 507–529.

3. Zhu, S., M.-Q. Yan, A. Masson, W. Chen, and Y.-P. Li. 2026. Cell signaling and transcriptional regulation of osteoclast lineage commitment, differentiation, bone resorption and diseases. Cell Discov. 12: 6.

4. Yin, L., C. Sun, J. Zhang, Y. Li, Y. Wang, L. Bai, and Z. Lei. 2025. Critical signaling pathways in osteoclast differentiation and bone resorption: mechanisms and therapeutic implications for periprosthetic osteolysis. Front. Cell Dev. Biol. 13: 1639430.

5. Coury, F., O. Peyruchaud, and I. Machuca-Gayet. 2019. Osteoimmunology of Bone Loss in Inflammatory Rheumatic Diseases. Front. Immunol. 10: 679.

6. Xu, F., and S. L. Teitelbaum. 2013. Osteoclasts: New Insights. Bone Res. 1: 11–26.

7. Boyce, B. F., and L. Xing. 2008. Functions of RANKL/RANK/OPG in bone modeling and remodeling. Arch. Biochem. Biophys. 473: 139–146.

8. Boyce, B. F., Y. Xiu, J. Li, L. Xing, and Z. Yao. 2015. NF-κB-Mediated Regulation of Osteoclastogenesis. Endocrinol. Metab. 30: 35–44.

9. Adamopoulos, I. E., and E. D. Mellins. 2015. Alternative pathways of osteoclastogenesis in inflammatory arthritis. Nat. Rev. Rheumatol. 11: 189–194.

10. Alippe, Y., and G. Mbalaviele. 2019. Omnipresence of inflammasome activities in inflammatory bone diseases. Semin. Immunopathol. 41: 607–618.

11. Tseng, H.-W., S. G. Samuel, K. Schroder, J.-P. Lévesque, and K. A. Alexander. 2022. Inflammasomes and the IL-1 Family in Bone Homeostasis and Disease. Curr. Osteoporos. Rep. 20: 170–185.

12. Schroder, K., and J. Tschopp. 2010. The inflammasomes. Cell 140: 821–832.

13. Wang, C., T. Yang, J. Xiao, C. Xu, Y. Alippe, K. Sun, T.-D. Kanneganti, J. B. Monahan, Y. Abu-Amer, J. Lieberman, and G. Mbalaviele. 2021. NLRP3 inflammasome activation triggers gasdermin D-independent inflammation. Sci. Immunol. 6: eabj3859.

14. Shin, H. J., I. S. Kim, J. K. Kim, and E.-K. Jo. 2026. Molecular mechanisms of NLRP3 inflammasome activation. Exp. Mol. Med. 58: 650–663.

15. Xia, S., Z. Zhang, V. G. Magupalli, J. L. Pablo, Y. Dong, S. M. Vora, L. Wang, T.-M. Fu, M. P. Jacobson, A. Greka, J. Lieberman, J. Ruan, and H. Wu. 2021. Gasdermin D pore structure reveals preferential release of mature interleukin-1. Nature 593: 607–611.

16. Kayagaki, N., O. S. Kornfeld, B. L. Lee, I. B. Stowe, K. O’Rourke, Q. Li, W. Sandoval, D. Yan, J. Kang, M. Xu, J. Zhang, W. P. Lee, B. S. McKenzie, G. Ulas, J. Payandeh, M. Roose-Girma, Z. Modrusan, R. Reja, M. Sagolla, J. D. Webster, V. Cho, T. D. Andrews, L. X. Morris, L. A. Miosge, C. C. Goodnow, E. M. Bertram, and V. M. Dixit. 2021. NINJ1 mediates plasma membrane rupture during lytic cell death. Nature 591: 131–136.

17. Broz, P. 2025. Pyroptosis: molecular mechanisms and roles in disease. Cell Res. 35: 334– 344.

18. Mbalaviele, G., and D. J. Veis. 2018. Inflammasomes in Bone Diseases. Experientia. Suppl. 108: 269–279.

19. Yu, C., C. Zhang, Z. Kuang, and Q. Zheng. 2021. The Role of NLRP3 Inflammasome Activities in Bone Diseases and Vascular Calcification. Inflammation 44: 434–449.

20. Li, Y., J. Ling, and Q. Jiang. 2021. Inflammasomes in Alveolar Bone Loss. Front. Immunol. 12: 691013.

21. Bauernfeind, F. G., G. Horvath, A. Stutz, E. S. Alnemri, K. MacDonald, D. Speert, T. Fernandes-Alnemri, J. Wu, B. G. Monks, K. A. Fitzgerald, V. Hornung, and E. Latz. 2009. Cutting edge: NF-kappaB activating pattern recognition and cytokine receptors license NLRP3 inflammasome activation by regulating NLRP3 expression. J. Immunol. 183: 787–791.

22. McKee, C. M., and R. C. Coll. 2020. NLRP3 inflammasome priming: A riddle wrapped in a mystery inside an enigma. J. Leukoc. Biol. 108: 937–952.

23. Molina-López, C., L. Hurtado-Navarro, C. J. García, D. Angosto-Bazarra, F. Vallejo, A. Tapia-Abellán, J. R. Marques-Soares, C. Vargas, S. Bujan-Rivas, F. A. Tomás-Barberán, J. I. Arostegui, and P. Pelegrin. 2024. Pathogenic NLRP3 mutants form constitutively active inflammasomes resulting in immune-metabolic limitation of IL-1β production. Nat. Commun. 15: 1096.

24. Cosson, C., R. Riou, D. Patoli, T. Niu, A. Rey, M. Groslambert, C. De Rosny, E. Chatre, O. Allatif, T. Henry, F. Venet, F. Milhavet, G. Boursier, A. Belot, Y. Jamilloux, E. Merlin, A. Duquesne, G. Grateau, L. Savey, A. T. Jacques Maria, A. Pagnier, S. Poutrel, O. Lambotte, C. Mallebranche, S. Ardois, O. Richer, I. Lemelle, F. Rieux-Laucat, B. Bader-Meunier, Z. Amoura, I. Melki, L. Cuisset, I. Touitou, M. Geyer, S. Georgin-Lavialle, and B. F. Py. 2024. Functional diversity of NLRP3 gain-of-function mutants associated with CAPS autoinflammation. J. Exp. Med. 221: e20231200.

25. 25. Zou, W., C. Wang, Y. Li, W. Jia, S. L. Teitelbaum, and G. Mbalaviele. 2025. Activation of the noncanonical inflammasome-GSDMD pathway triggers pyroptosis in bone marrow and promotes periosteal bone formation. .

26. Bonar, S. L., S. D. Brydges, J. L. Mueller, M. D. McGeough, C. Pena, D. Chen, S. K. Grimston, C. L. Hickman-Brecks, S. Ravindran, A. McAlinden, D. V. Novack, D. L. Kastner, R. Civitelli, H. M. Hoffman, and G. Mbalaviele. 2012. Constitutively activated NLRP3 inflammasome causes inflammation and abnormal skeletal development in mice. PloS One 7: e35979.

27. Qu, C., S. L. Bonar, C. L. Hickman-Brecks, S. Abu-Amer, M. D. McGeough, C. A. Peña, L. Broderick, C. Yang, S. K. Grimston, J. Kading, Y. Abu-Amer, D. V. Novack, H. M. Hoffman, R. Civitelli, and G. Mbalaviele. 2015. NLRP3 mediates osteolysis through inflammation-dependent and -independent mechanisms. FASEB J. Off. Publ. Fed. Am. Soc. Exp. Biol. 29: 1269–1279.

28. Xiao, J., C. Wang, J.-C. Yao, Y. Alippe, T. Yang, D. Kress, K. Sun, K. L. Kostecki, J. B. Monahan, D. J. Veis, Y. Abu-Amer, D. C. Link, and G. Mbalaviele. 2020. Radiation causes tissue damage by dysregulating inflammasome-gasdermin D signaling in both host and transplanted cells. PLoS Biol. 18: e3000807.

29. Kaur, K., Y. Alippe, C. Wang, N. P. Semenkovich, M. G. Hassan, S. Bhagat, K. Khanna, Y. Li, N. Pokhrel, T. Peterson, E. L. Scheller, D. J. Veis, Y. Abu-Amer, R. Faccio, and G. Mbalaviele. 2026. Activation of the NLRP3 inflammasome in osteoclasts is suppressed by a Tmem178- dependent mechanism that restricts Ca^2+^ influx. Sci. Signal. 19: eaea2753.

30. Alippe, Y., D. Kress, B. Ricci, K. Sun, T. Yang, C. Wang, J. Xiao, Y. Abu-Amer, and G. Mbalaviele. 2021. Actions of the NLRP3 and NLRC4 inflammasomes overlap in bone resorption. FASEB J. Off. Publ. Fed. Am. Soc. Exp. Biol. 35: e21837.

31. Otero, J. E., T. Chen, K. Zhang, and Y. Abu-Amer. 2012. Constitutively active canonical NF- κB pathway induces severe bone loss in mice. PloS One 7: e38694.

32. Schmacke, N. A., F. O’Duill, M. M. Gaidt, I. Szymanska, J. M. Kamper, J. L. Schmid-Burgk, S. C. Mädler, T. Mackens-Kiani, T. Kozaki, D. Chauhan, D. Nagl, C. A. Stafford, H. Harz, A. L. Fröhlich, F. Pinci, F. Ginhoux, R. Beckmann, M. Mann, H. Leonhardt, and V. Hornung. 2022. IKKβ primes inflammasome formation by recruiting NLRP3 to the trans-Golgi network. Immunity 55: 2271–2284.e7.

33. Nanda, S. K., A. R. Prescott, C. Figueras-Vadillo, and P. Cohen. 2021. IKKβ is required for the formation of the NLRP3 inflammasome. EMBO Rep. 22: e50743.

34. Mehrotra, P., S. Maschalidi, L. Boeckaerts, C. Maueröder, R. Tixeira, J. Pinney, J. Burgoa Cardás, V. Sukhov, Y. Incik, C. J. Anderson, B. Hu, B. N. Keçeli, A. Goncalves, L. Vande Walle, N. Van Opdenbosch, A. Sergushichev, E. Hoste, U. Jain, M. Lamkanfi, and K. S. Ravichandran. 2024. Oxylipins and metabolites from pyroptotic cells act as promoters of tissue repair. Nature 631: 207–215.

35. Phulphagar, K., L. I. Kühn, S. Ebner, A. Frauenstein, J. J. Swietlik, J. Rieckmann, and F. Meissner. 2021. Proteomics reveals distinct mechanisms regulating the release of cytokines and alarmins during pyroptosis. Cell Rep. 34: 108826.

36. Wright, S. S., P. Kumari, V. Fraile-Ágreda, C. Wang, S. Shivcharan, S. Kappelhoff, E. G. Margheritis, A. Matz, S. O. Vasudevan, I. Rubio, M. Bauer, B. Zhou, S. K. Vanaja, K. Cosentino, J. Ruan, and V. A. Rathinam. 2025. Transplantation of gasdermin pores by extracellular vesicles propagates pyroptosis to bystander cells. Cell 188: 280–291.e17.

37. Alippe, Y., C. Wang, B. Ricci, J. Xiao, C. Qu, W. Zou, D. V. Novack, Y. Abu-Amer, R. Civitelli, and G. Mbalaviele. 2017. Bone matrix components activate the NLRP3 inflammasome and promote osteoclast differentiation. Sci. Rep. 7: 6630.

38. Ruscitti, P., P. Cipriani, F. Carubbi, V. Liakouli, F. Zazzeroni, P. Di Benedetto, O. Berardicurti, E. Alesse, and R. Giacomelli. 2015. The role of IL-1β in the bone loss during rheumatic diseases. Mediators Inflamm. 2015: 782382.

39. Chen, T., L. Jin, J. Li, and Y. Liu. 2024. Pyroptosis mediates osteoporosis via the inflammation immune microenvironment. Front. Immunol. 15: 1371463.

