## Supplemental figures 1-5 for "Pyroptotic cell-derived factors promote osteoclast differentiation"

Figure S1

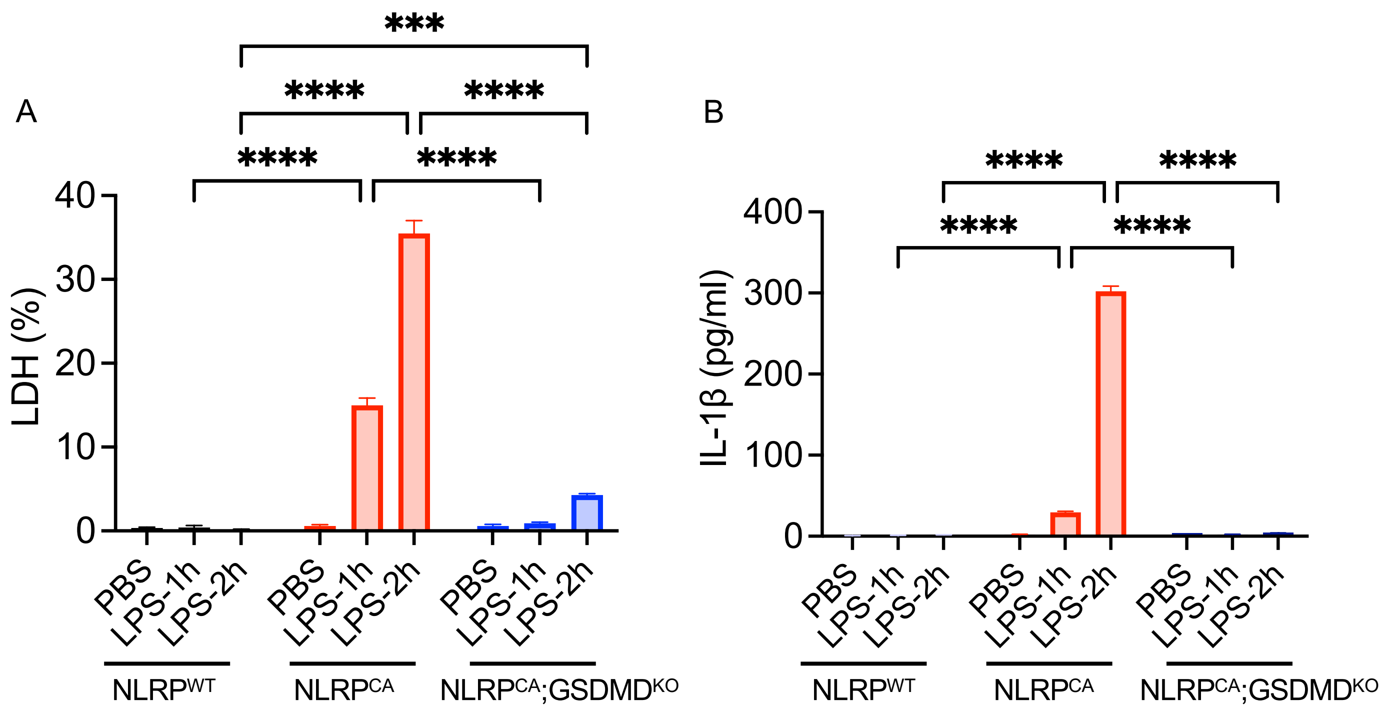


**Figure S1. LPS alone causes pyroptosis in NLRP3$^{CA}$, but not in NLRP3$^{WT}$ or NLRP3$^{CA}$;GSDMD$^{KO}$ BMDMs**

BMDMs isolated from NLRP3$^{WT}$, NLRP3$^{CA}$ and NLRP3$^{CA}$;GSDMD$^{KO}$ mice were treated with PBS or 100 ng/ml LPS for 1 h or 2 h. KO, knockout. Levels of LDH (A) and IL-1β were measured in CM (B). Samples were run in triplicates and data are presented as mean ± SD. ***P<.001, ****P<.0001, two-way ANOVA test.

Figure S2

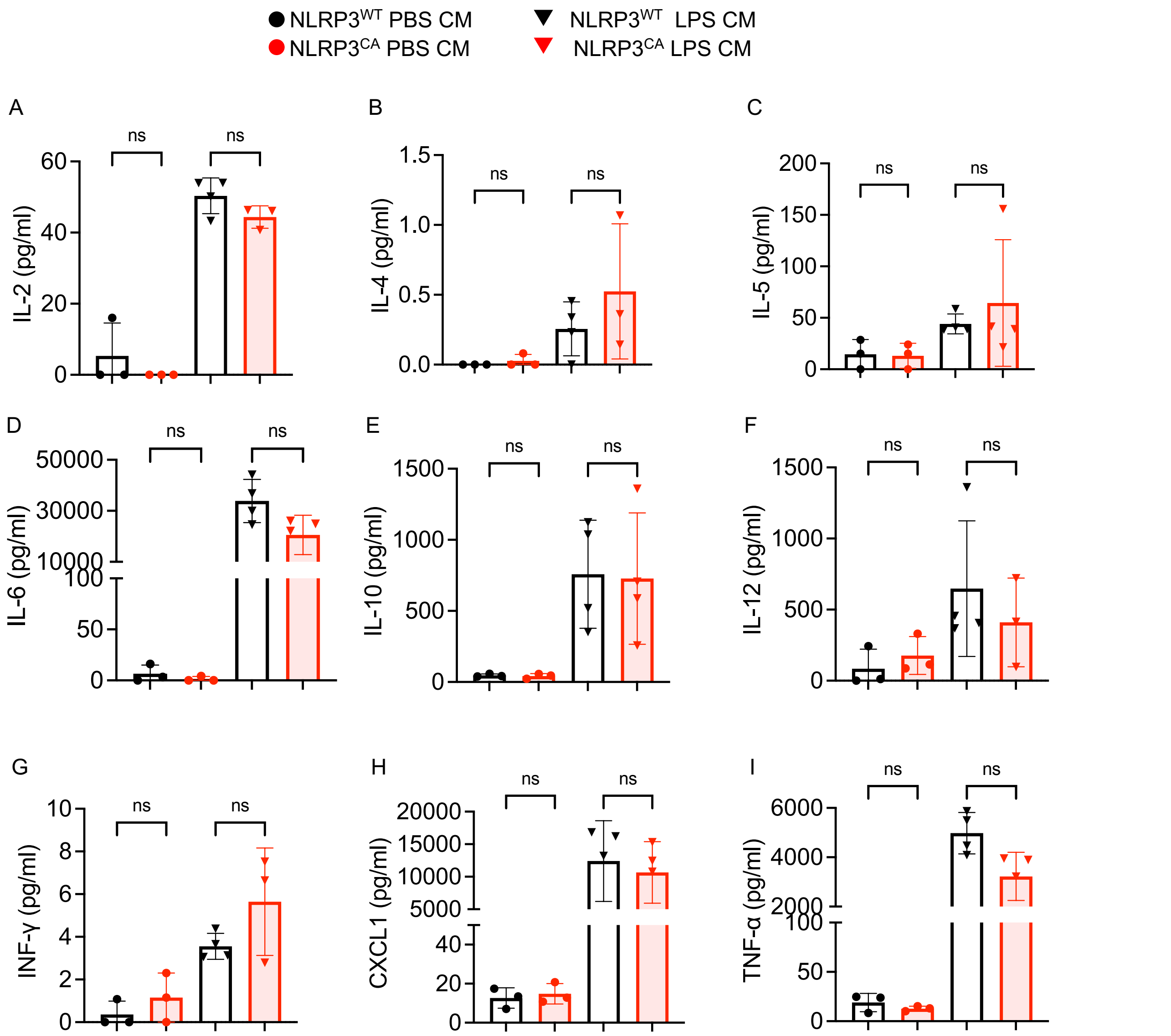


**Figure S2. LPS does not induce generalized inflammation through NLRP3CA**

The levels of IL-2 (A), IL-4 (B), IL-5 (C), IL-6 (D), IL-10 (E), IL-12 (F), IFN-ɣ (G), CXCR1 (H) and TNF-α (I) were measured in NLRP3WT PBS CM, NLRP3CA PBS CM, NLRP3WT LPS CM. and NLRP3CA LPS CM (n=3~4). Data are presented as mean ± SD. ***P<.001, ****P<.0001; one-way ANOVA test.

Figure S3

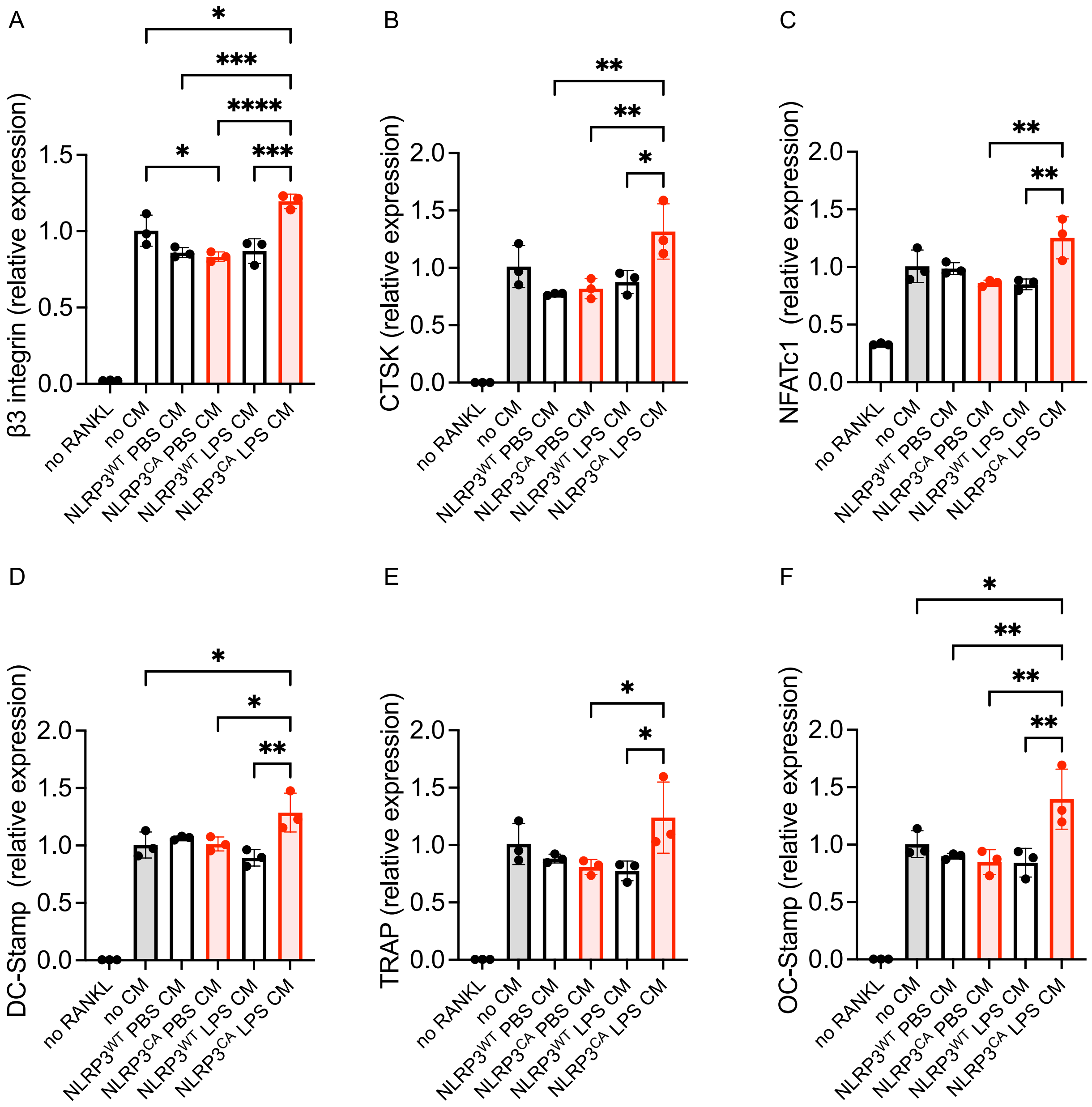


**Figure S3. NLRP3$^{CA}$ LPS CM induces stronger expression of OC markers compared to NLRP3$^{WT}$ LPS CM**

NLRP3$^{WT}$ PBS CM, NLRP3$^{CA}$ PBS CM, NLRP3$^{WT}$ LPS CM, and NLRP3$^{CA}$ LPS CM were added to WT BMDMs primed with RANKL for 2 days. Cells were cultured for additional 2-3 days in the presence of RANKL and CM refreshed every day. Gene expression was analyzed by qPCR. Samples were run in triplicates and data are presented as mean ± SD. *P<.05, **P<.01, ***P<.001, ****P<.0001; one-way ANOVA test.

Figure S4

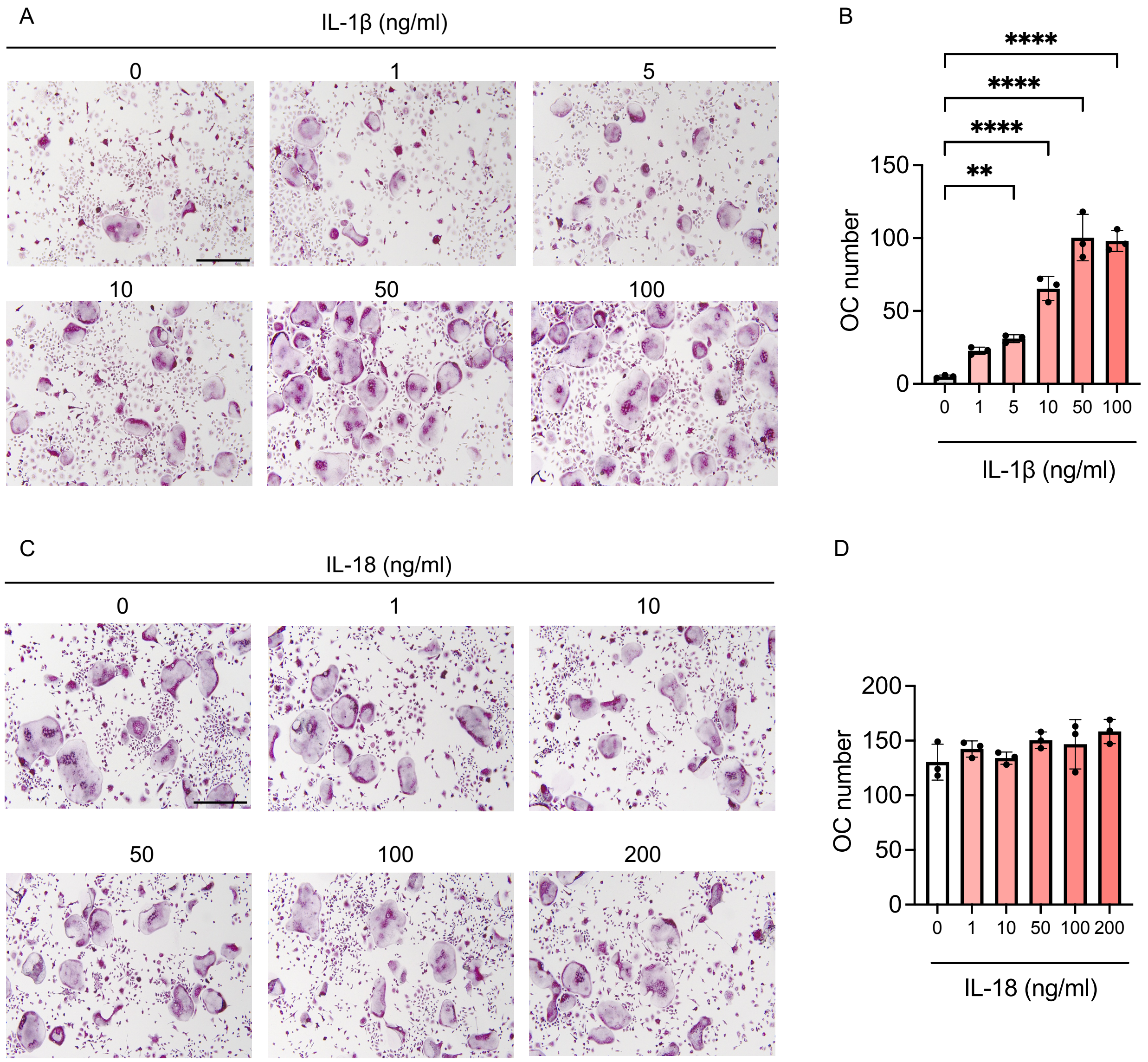


**Figure S4. IL-1β but not IL-18 enhances OC differentiation induced by RANKL**

IL-1β or IL-18 added to WT BMDMs primed with RANKL for 2 days. Fresh IL-1β or IL-18 was added daily to RANKL-exposed cultures. Cells were stained for TRAP after mature osteoclasts had formed. Representative TRAP-stained images of cells treated with IL-1β (A) or IL-18 (C) in different concentrations, together with quantification of osteoclast numbers (B, D), are shown (n=3). Data are presented as mean ± SD. ***P<.001, ****P<.0001; one-way ANOVA test.

Figure S5

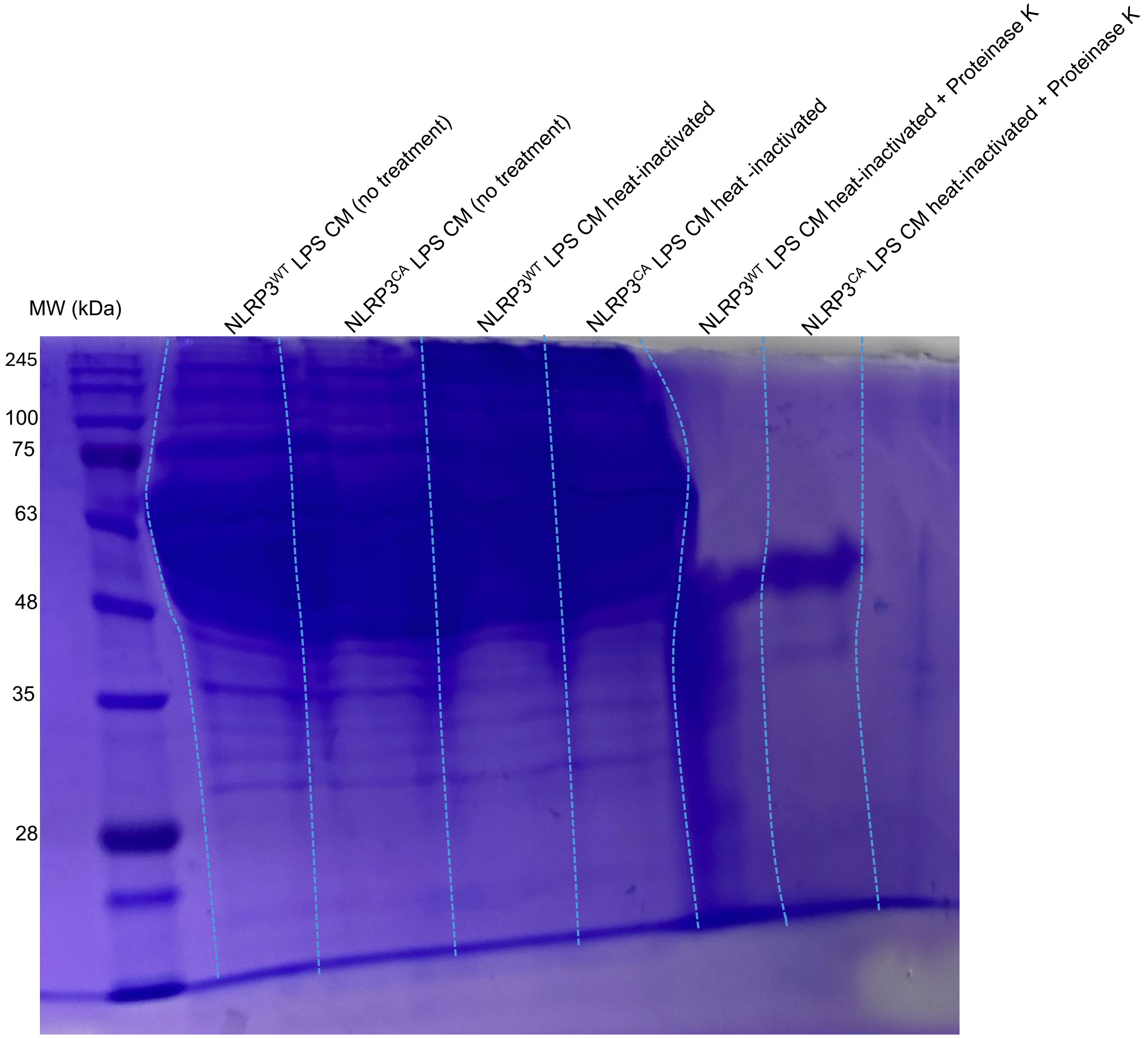


**Figure S5. Inactivation of proteins by the heat in the absence or presence of proteinase K**

Five µl from untreated $NLRP3^{WT}$ LPS CM, $NLRP3^{WT}$ LPS CM (heat inactivated), $NLRP3^{WT}$ LPS CM (heat inactivated + proteinase K), untreated $NLRP3^{CA}$ LPS CM, $NLRP3^{CA}$ LPS CM (heat inactivated) and $NLRP3^{CA}$ LPS CM (heat inactivated + proteinase K) was subjected to gel electrophoresis, and the gel was stained with Coomassie Brilliant Blue.

Table S1. List of primers for qPCR

| Primers | Fw sequence | Rv sequence |
|---|---|---|
| Cyclophilin B | AGCATACAGGTCCTGGCATC | TTCACCTTCCCAAAGACCAC |
| β3 intergrin | TTCGACTACGGCCAGATGATT | GGAGAAAGACAGGTCCATCAAGT |
| OC stamp | CTGTAACGAACTACTGACCCAGC | CCCAGGCTTAGGAAGACGAAG |
| DC stamp | ACAAAGCAACAGACTCCCAAAT | GGGGACTTATGTGTTTCCACG |
| CTSK | AGGCAGCTAAATGCAGAGGGTACA | AGCTTGCATCGATGGACACAGAGA |
| NFATc1 | CCCGTCACATTCTGGTCCAT | CAAGTAACCGTGTAGCTGCACAA |
| TRAP | CGTCTCTGCACAGATTGCAT | AAGCGCAAACGGTAGTAAGG |
